# Variational inference accelerates accurate DNA mixture deconvolution

**DOI:** 10.1101/2022.12.01.518640

**Authors:** Mateusz Susik, Ivo F. Sbalzarini

## Abstract

We investigate a class of DNA mixture deconvolution algorithms based on variational inference, and we show that this can significantly reduce computational runtimes with little or no effect on the accuracy and precision of the result. In particular, we consider Stein Variational Gradient Descent (SVGD) and Variational Inference (VI) with an evidence lower-bound objective. Both provide alternatives to the commonly used Markov-Chain Monte-Carlo methods for estimating the model posterior in Bayesian probabilistic genotyping. We demonstrate that both SVGD and VI significantly reduce computational costs over the current state of the art. Importantly, VI does so without sacrificing precision or accuracy, presenting an overall improvement over previously published methods.

## 1 Introduction

DNA mixture analysis using probabilistic genotyping (PG) software is at the core of forensic science methodologies. While the greatest attention has to be placed on the accuracy of the results, there are other factors that play a role. Recently, it has been shown how the precision of a PG system can be improved by reducing run-to-run variability [15]. Another factor that can be considered is the computational runtime. In many cases, estimating a likelihood ratio (LR) can take more than an hour. Given the workload forensic laboratories face, faster PG algorithms are desirable as long as they produce results of equal accuracy and precision.

The accuracy of a PG result directly depends on how well the PG algorithm is able to approximate or estimate the probabilities of the parameters of a model given an observed electropherogram. Often, this is formulated as a Bayesian estimation problem where the task is to estimate the posterior distribution. In state-of-the-art PG models [15, 16], it is not possible to calculate the posterior directly by integrating over the parameters in a reasonable timeframe. Instead, sampling algorithms are used, most prominently Markov-Chain Monte-Carlo (MCMC) methods with random walk [16] or Hamiltonian proposal distributions [15]. These algorithms then estimate the unknown posterior distribution from the electropherogram data. The posterior distribution lives in a space whose dimensionality depends on the number of unknown parameters to be estimated. Since the computational cost of a PG algorithm scales with the number of model evaluations it requires, it is desirable to have PG methods that estimate the posterior distribution as accurately as possible using as few samples as possible.

The cost-performance trade-off is a classic research topic in Bayesian inference [18]. There, besides MCMC methods, also other types of algorithms have been proposed. In particular variational methods have been successful at improving the computational performance of Bayesian inference [3]. It therefore seems natural to also adopt these approaches in DNA mixture deconvolution and benchmark their performance against the state of the art in PG.

Here, we present implementations of two variational inference techniques adapted to PG applications: variational inference with an evidence lower bound objective (VI) [3] and Stein Variational Gradient Descent (SVGD) [11]. We show that both SVGD and VI achieve shorter runtimes than the MCMC-based methods. Importantly, VI does so without sacrificing precision or accuracy, presenting an overall improvement over the state of the art in PG.

## 2 Materials and methods

In Bayesian inference, the main task is to maximise the posterior probability of a model *M* given data *V* :

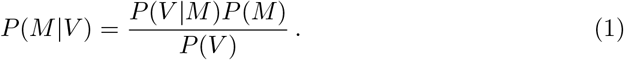

The evidence *P*(*V*) is a constant that is independent of the model. Since the location of the maximum in the posterior does not depend on this normalisation, it is often neglected. In the context of PG, namely to obtain the weights of the genotype sets for LR calculation, this is also the case because the evidence crosses out when computing the ratio of likelihoods.

Therefore, a Bayesian inference algorithm can in practice be seen as consisting of two parts: an estimator and a model (Figure 1). The model defines the unnormalised posterior, and the estimator defines the way how an approximation of this distribution is obtained. These two parts are largely independent of each other, meaning that, for example, an estimator can be replaced with another one.

**Figure 1.**
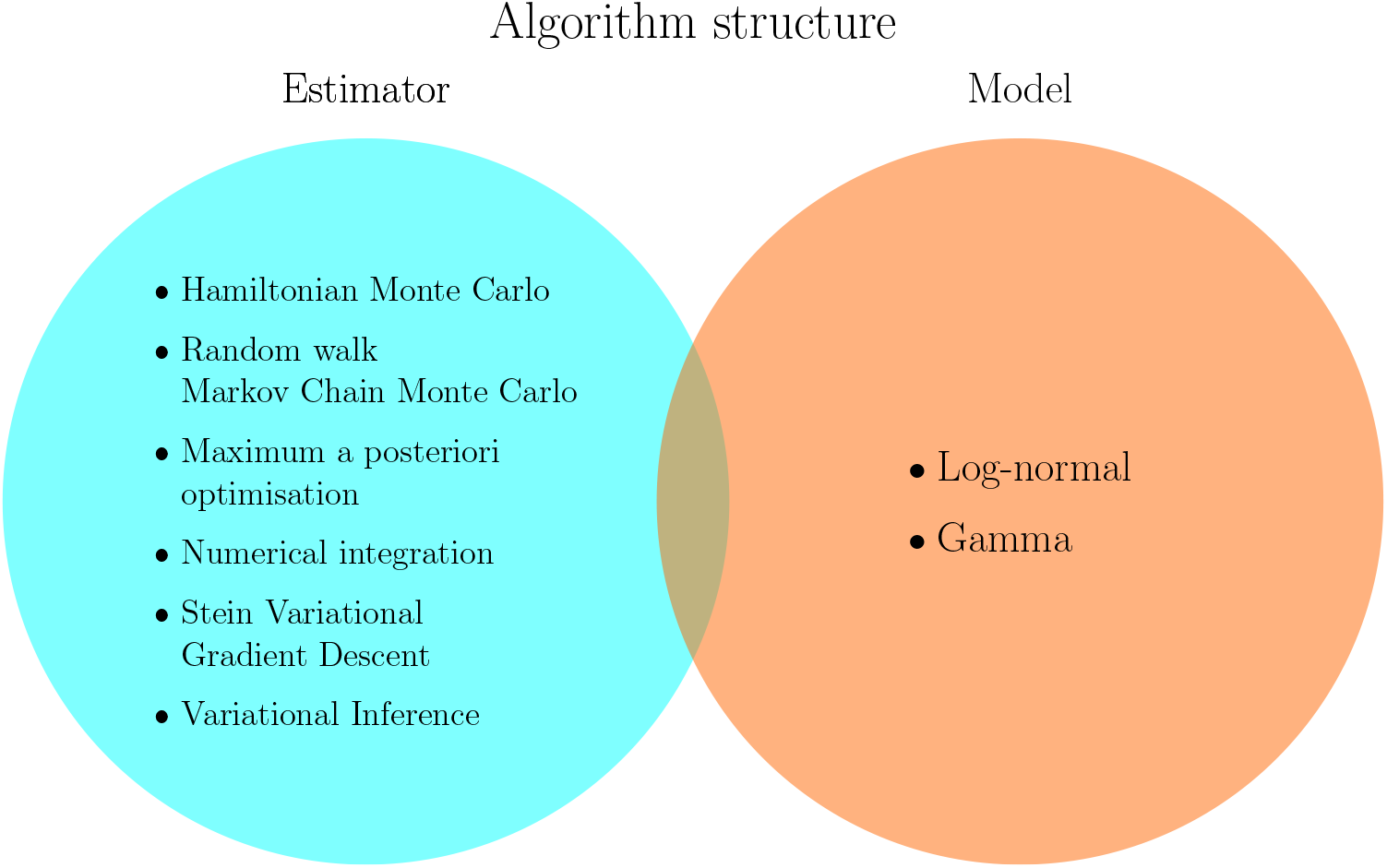
The basic structure of a probabilistic genotyping algorithm: The algorithm consists of an *estimator*, a tool for approximating a distribution, and a *model* that defines the unnormalised posterior, usually by defining the likelihood and the prior. Different popular choices of estimator methods and PG models are given in the circles. Their choices are largely mutually independent.

In practice, this ideally means that data scientists can create a model based on observed data and/or theoretical knowledge, while different estimators can be used interchangeably in order to optimise for computational cost and/or accuracy and precision of the PG result. Yet, the model might constrain the choice of the estimator, as different estimators have different structural limitations.

### 2.1 Model definition

In this work, we follow the model and hyper-parameterisation used in Hamiltonian Monte Carlo (HMC) [15] with a few exceptions as described next. These minor adaptations are required for variational estimators to work correctly. First, consider the likelihood probability distribution suggested by the authors of STRmix™ [16]:

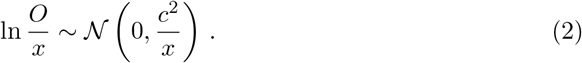

This postulates that the ratio of the observed peak height *O* to the expected peak height *x* follows a log-normal distribution with mean 0 and a variance proportional to the square of a parameter *c* simulated by the model and inversely proportional to the expected peak height *x*. During inference, the estimator might thus try values of *c* close to 0, as long as the Gamma prior [8] does not forbid this. The probability density *f*_LogN_ of this parameterised log-normal distribution can then reach arbitrarily large values:

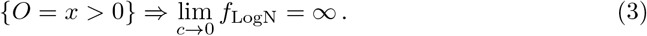

This is the first issue hampering the use of variational inference methods.

The second issue is similar: Consider the prior probability distributions for the locus-specific amplification efficiencies (LSAE) *α* with a hyper-parameter *σ*_*α*_ set by the laboratory:

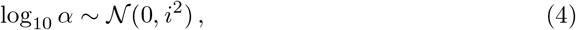

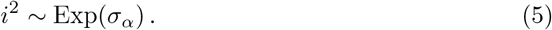

The prior density for the LSAEs *f*_LSAE_ is then also unbounded:

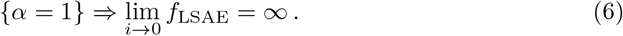

These singularities are not a problem for HMC estimators, who will avoid them because of the high curvature of the posterior in the vicinity of the singularities. When the sampler tries to explore these parts of the posterior, the trajectory of the simulated Hamiltonian differs too much from the expected Hamiltonian. The sample is then rejected and marked as a “divergence”. These samples then negatively impact the runtime of the algorithm. The convergence of the chains is slower and, in extreme cases, can lead to a bias in the estimate.

Variational inference estimators, however, are not able to work with posteriors that contain singularities. We therefore modify the model as follows:

- use shifted log-normal (LogN) priors for allele and stutter peak heights, instead of the Gamma priors used by STRmix™.
- use a log-logistic (LogL) hyper-prior for the LSAE standard deviation, instead of an exponential one on LSAE variance (Eq. 5).

The exact choice of the prior distribution families, in our case log-normal and log-logistic, is not crucial as long as the estimates densities protect the algorithms from the singularities.

To estimate the parameters of these prior distributions, we start with a prior-less posterior estimation of a training set of DNA mixtures. We then extract, for each mixture, the samples (peak height variance, stutter height variance, LSAE variance) that are larger than the 90th percentile of the estimated parameter distribution. Finally, we choose the parameters of the prior distributions that maximise the likelihood of this subsample.

For the following results, we used 37 2- and 3-contributor filtered Globalfiler™ mixtures that were not a part of the test benchmark [12, 14]. For the list of mixtures, please see Supplementary Material 1. The estimated priors are presented in Table 1 and compared with the priors from Riman et al. [12] in Figure 2.

**Table 1.** Estimated priors based on 37 2- and 3-contributor Globalfiler™ ProvedIT mixtures not used for the test benchmark.

| Prior | Distribution |
| --- | --- |
| stutter peak height standard deviation | $c_s - 6.086 \sim \text{LogN}(2.485, 1.031)$ |
| allelic peak height standard deviation | $c_p - 1.63 \sim \text{LogN}(1.975, 0.325)$ |
| LSAE standard deviation hyper-prior | $i \sim \text{LogL}(-1.894, 0.236)$ |

**Figure 2.**
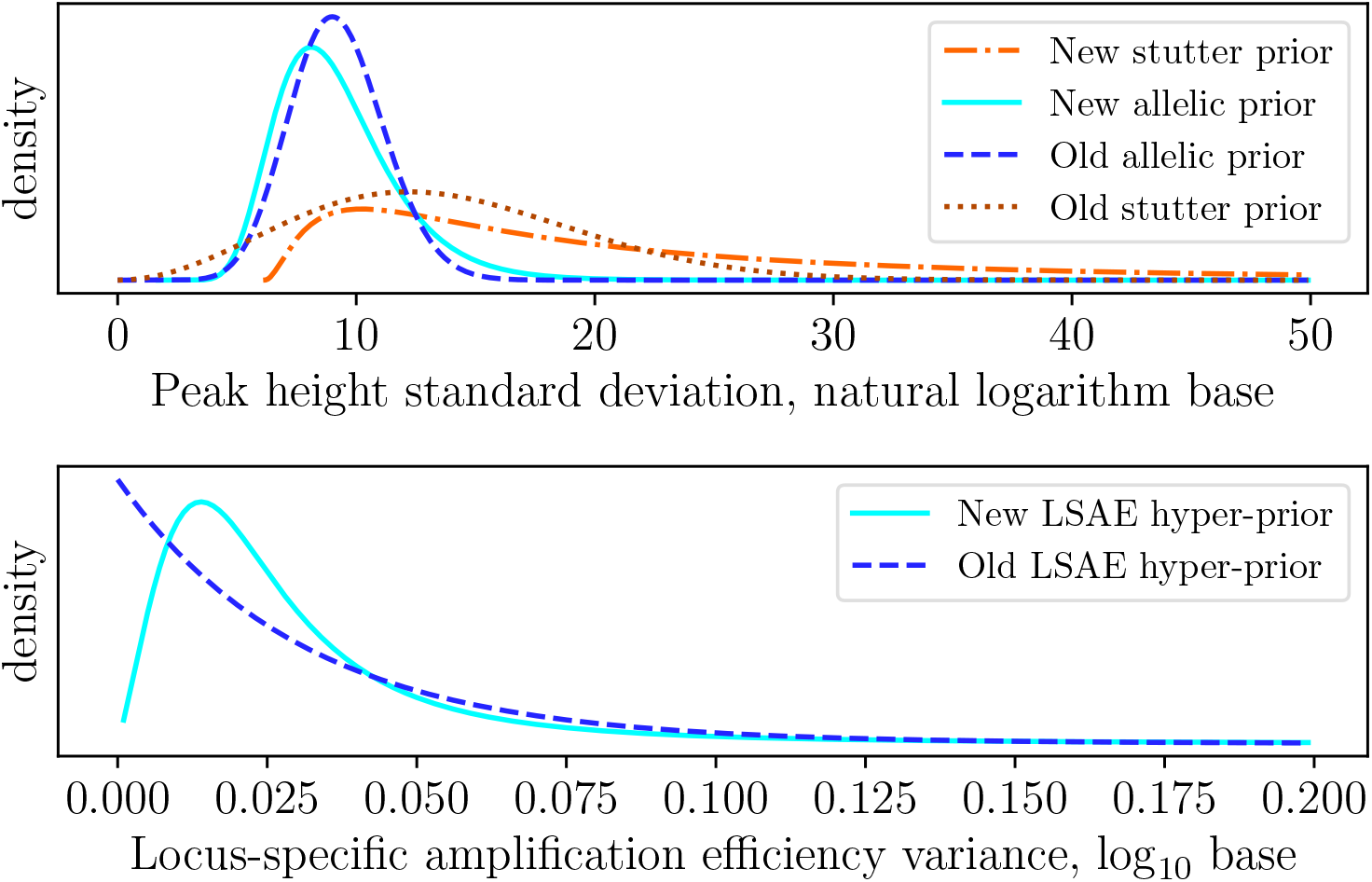
Comparison of the singularity-free priors estimated using our method with those from previous studies [12, 15]. Peak height standard deviation is expressed in natural logarithms, i.e. ln 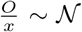, whereas LSAE variance is presented in a decimallogarithms base: log_10_

The shifted peak-height standard deviation priors prohibit parameter values smaller than the shift. One could argue that this introduces a bias into the analysis, as the “true” value of the parameter might not be accessible to the estimator. While this is true, it is not specific to our work. Indeed, also in other models these distributions are shifted to the right of the estimates from the model, see, e.g., Figure 4 from Taylor et al. [17]. Moreover, a natural meaning of the peak height standard deviation parameter is the confidence of the model when estimating peak heights. The lower the value of the parameter, the more confident the model. An overestimation of these parameters might be then desirable, since this increases the level of uncertainty of the estimator.

**Figure 3.**
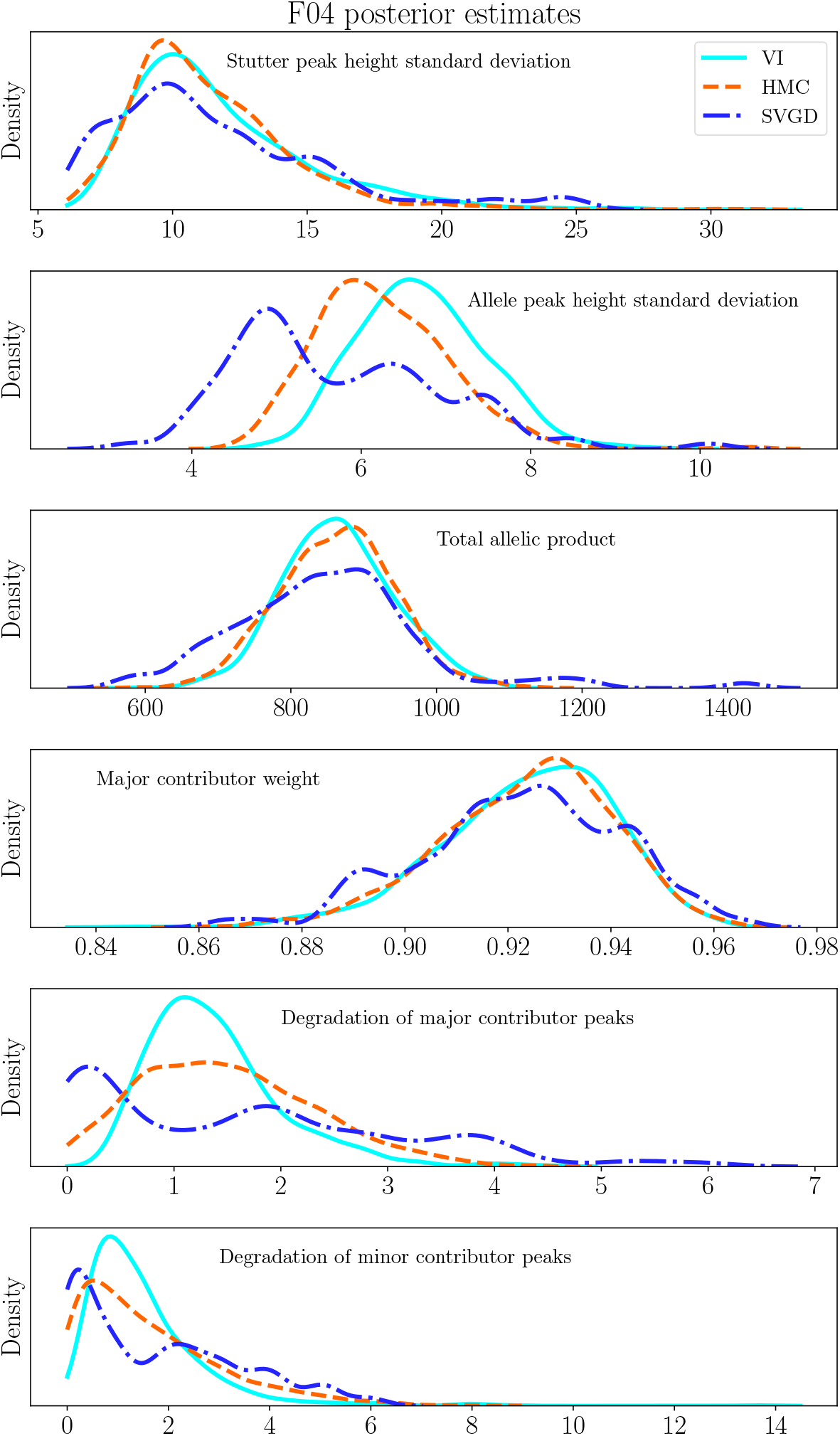
Comparison of the posteriors estimated for the F05 ProvedIT mixture by different algorithms. We plot the marginal densities of the posteriors for the parameters: peak-height standard deviations, total allelic product, major contributor weight, and contributors’ template degradation. All plots are created using kernel density estimation. The estimation has been performed on 1000 samples from the Gaussian estimated by VI, 800 samples from HMC, and 100 particles from SVGD. Given the smaller sample size from SVGD, we decreased the kernel bandwidth two-fold, resulting in plots that are less smooth for SVGD.

**Figure 4.**
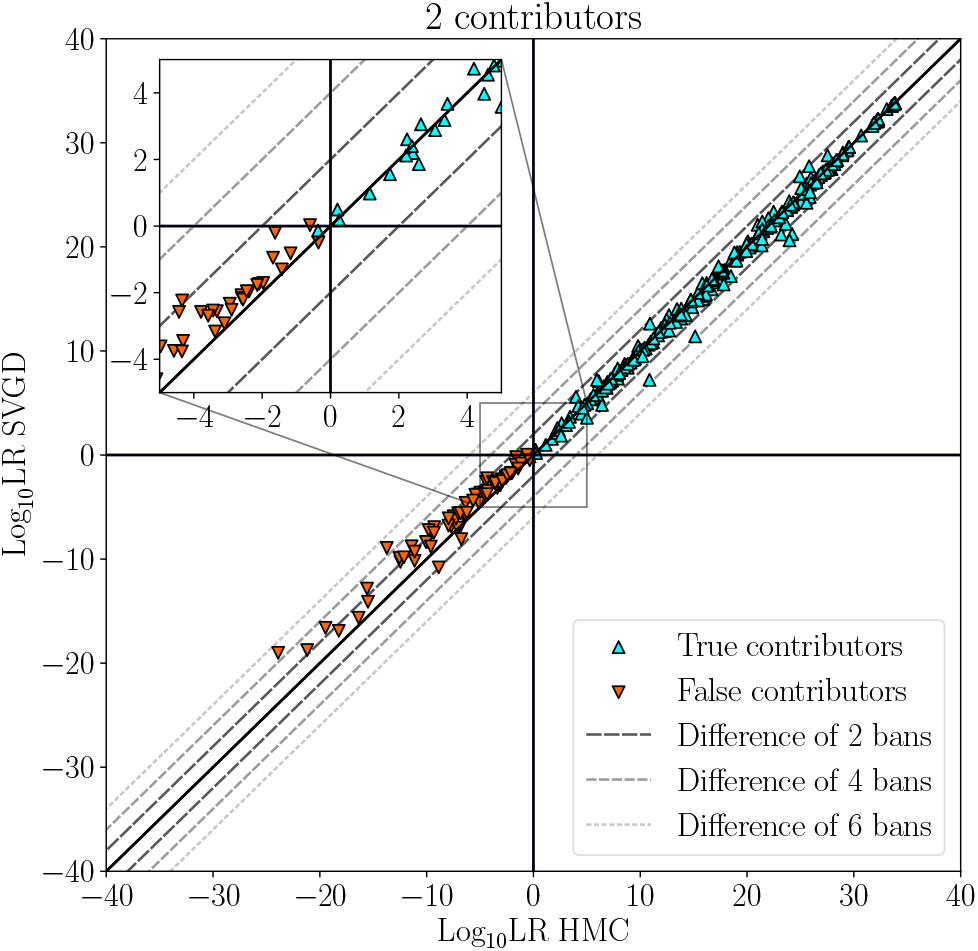
Comparison of results obtained by SVGD and HMC on 2-contributor mixtures.

### 2.2 Variational inference

In Bayesian inference, one typically distinguishes sampling methods, such as MCMC and HMC, from variational methods. Sampling methods iteratively draw samples from the posterior. They construct a Markov chain of the samples. The chains, as long as they are ergodic, converge to the desired stationary distribution, which is the (unnormalized) posterior of the model in the limit of infinitely many samples. In practice, however, the number of samples is finite. Therefore, convergence criteria (e.g. the Gelman-Rubin test) are used to determine when to stop the sampler. This inherently trades off computational cost, which is proportional to the number of samples drawn, with accuracy and precision (i.e., reproducibility) of the result. Variational methods avoid this trade-off by directly building an approximation to the posterior. This approximation comes from a family of distributions, called the variational family, which is selected by the design of the algorithm. For simpler models, it is sometimes possible to construct an exact variational family, but this is not the case for the model considered here.

#### 2.2.1 Variational inference with an evidence lower-bound objective

The first variational method we consider uses multivariate normal distributions as the variational family *Q*(*M*). It then aims to minimise the Kullback-Leibler (KL) divergence:

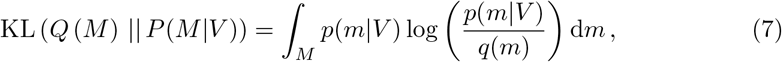

where *p*(*m*|*V*) and *q*(*m*) are the densities of the posterior and the variational distributions, respectively. The final result of VI is the distribution from the variational family that resembles the posterior the most, where resemblance is measured by the KL divergence between the two distributions. While KL divergence cannot be calculated directly in our case, it can be shown that in general:

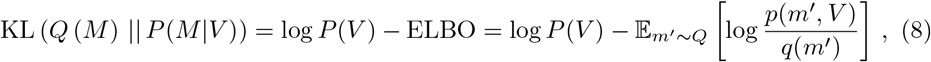

where ELBO is the so-called “evidence lower-bound objective”. Since the log-evidence is a constant independent of the model, the KL divergence is minimised by maximising ELBO. This formulation of the optimisation problem is flexible w.r.t. the choice of the variational family. In order to consider a wide choice of possible solutions, we here use a multivariate Gaussian family with full covariance matrix.

This choice of variational family could harm the quality of the results if the true posterior distribution cannot be approximated by multivariate Gaussians. Empirically, however, we observe that the posterior estimated by HMC, which should be a close approximation of the true posterior, is similar to a transformed multivariate Gaussian. We therefore expect VI to work well. We compare the posterior estimated by the different methods in Figure 3. We run all three methods identically with the priors presented in Section 2.1 on the F04 RD14−0003−42 43−1;9−M3a−0.15GF−Q0.5 06.15sec (referred to as “F04”) mixture from the ProvedIT dataset [1]. Since the distributions are transformed [15], the marginal density plots do not always look Gaussian. A transformed normal distribution output by VI, however, approximates well the answer provided by HMC. Still, we note some differences exist, such as decreased variance of the estimated marginal degradation distributions.

We maximise the ELBO using the Adamax algorithm [11]. This algorithm internally uses a Monte-Carlo estimator for the ELBO according to Eq. 8. We use 10 samples per estimation and a variable learning rate (LRate) schedule: The first iterations are performed with LRate=0.01; then, after every 100 iterations, the LRate is multiplied by 1.5 until it reaches 0.05. This adaptation prevents diverging gradients that could otherwise be caused by bad initialisation of the variational family. Indeed, during the first few iterations, gradient computation could become numerically unstable, as an outlier sampled from the variational distribution could cause it to significantly depart from reasonable parameter values. Convergence of the optimiser is monitored by comparing the mean value of the ELBO every 100 iterations. If this mean is smaller than the mean from the previous hundred iterations, the optimiser is stopped, since no further improvement was achieved.

#### 2.2.2 Stein variational gradient descent

Stein variational gradient descent (SVGD) [11] intends to find a composition of transformations for an ensemble of *n* particles in the *d*-dimensional parameter space with positions 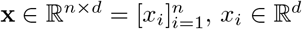, that maps an initial distribution to the best approximation of the true posterior as quantified by the KL divergence. Each individual transformation is defined as:

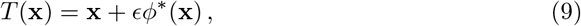

where *ϵ* is a step size and *ϕ*^∗^ : ℝ^*n*×*d*^ ↦ ℝ^*n*×*d*^ is a map defining a per-particle direction vector that maximises the rate of decrease of the KL divergence between the current transformed variational distribution *q*_[*T*]_ and the unnormalised posterior *p*:

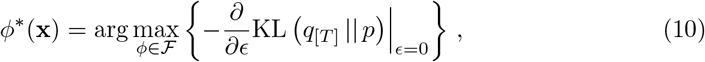

where *T* (*x*) ∼ *q*_[*T*]_ and *x* ∼ *q*. The transformation for each particle *x*_*i*_ depends on the other particles within some neighborhood. The authors of the method take the function space of the transformations to be the unit ball in a vector-valued reproducing kernel Hilbert space (RKHS) with positive-definite kernel *k*(·,·). Theorem 3.1 from Liu & Wang [11] tells us that the rate of decrease can be expressed in closed form as an expectation:

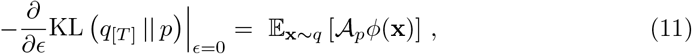

Where

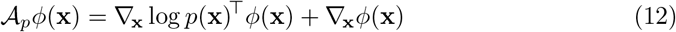

is the *Stein operator* [13]. The maximum in Eq. 10 is called the *Stein discrepancy*.

It has been shown that when ℱ is the unit ball in the RKHS, then the Stein discrepancy is a well-defined expectation value [11]:

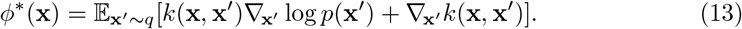

Intuitively, Eq. 13 means that:

- The particles prefer to be in regions of high probability density, as indicated by the gradient of the log-posterior. The log-posterior of the neighbours will dominate this term for any particle given the multiplication with the kernel.
- At the same time they are repelled from one another by the gradient of the kernel in order to not collapse into a single mode and cover the whole posterior.

To provide a computable update rule, the expectation is estimated by the average over the set of particles:

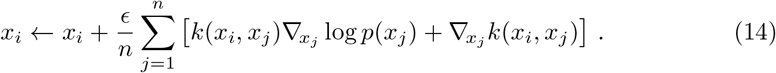

Note that Equation 11 holds for *ϵ* = 0, meaning that if we wanted to follow the exact optimal trajectory, we would have to use continuous updates. Any computable version must therefore discretise the update trajectories using a finite step size *ϵ*. As long as this step size is small enough, the particles still represent a sample from the correct posterior after a sufficient number of transformations, as demonstrated by Liu & Wang [11]. Following Liu & Wang [11], we use a radial basis function kernel 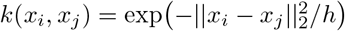, and we set bandwidth *h* to med^2^/ln *n* where med is the median of the pairwise distances between the *n* particles. In our experiments, we use 100 particles and run 500 updates. Our implementation uses the Adamax algorithm [9] with learning rate 0.25 to efficiently approximate the gradient descent in Eq. 14. Adjusting the learning rate, as done for VI, is not necessary, since instabilities in the gradient computation cannot occur.

When only one particle is used, *n* = 1, SVGD reduces to maximum a posteriori estimation, which is the method used in Euroformix [19].

We reduce memory consumption by iteratively evaluating the unnormalised posterior *p* in 10 equal batches of 10 particles each. This hyper-parameter can be adjusted depending on the available computational resources. Memory consumption scales linearly with batch size.

### 2.3 Implementation details

From the estimated posteriors, we compute LRs as described [15]. We first draw a sample from the estimated posterior distribution. For SVGD, the particles directly represent the sample [11]. For VI, we sample 1000 points from the estimated Gaussian. We note that increasing the number of particles in SVGD would linearly increase the computational time, while in case of VI we sample from an explicit Gaussian distribution only after the inference is finished, and thus we can afford a larger sample. Subsequently, the same approach as in HMC is used for both algorithms: we calculate the deconvolution by averaging the values across all samples.

We implement both the SVGD and VI estimators using the Tensorflow Probability library [6]. Gradients are computed using Tensorflow’s automatic differentiation. All benchmarks are performed on NC4as T4 v3 Azure cloud GPU instances unless specified otherwise.

## 3 Results

We validate the use of variational inference for forensic PG and provide a comparative study between the two presented methods, SVGD and VI, and HMC. We benchmark three characteristics important for any PG system: the accuracy of the method, it’s precision, and the computational runtime. We use the term *scenario* to indicate the combination of a DNA mixture with a certain prosecutor/defendant hypothesis.

### 3.1 Accuracy

In order to quantify the accuracy of the methods, we compare their results on the ProvedIT mixtures from the NIST comparative study [12]. We observe almost identical performance in terms of the ROC area-under-the-curve (AUC) between HMC (0.99896) and VI (0.99887), see Table 2. SVGD performs slightly worse (0.99843), which is caused by one scenario with true contributor #33 resulting in a large negative log_10_ LR of -16.3753, see Figure 6. This is mixture E05, which was already described in Supplementary Material 2 of Ref. [14]. The LR for the locus D12S391 and the sub-sub-source hypothesis that resulted in the largest LR overall is 2.71 · 10^−27^. When we consider the same alternative scenario as in Ref. [14], with the peak 18.3 (1220 RFU) added in locus D12S391, SVGD estimates a log_10_ LR of 11.646, which is close to the HMC estimate of 12.8367 in the same case. The sub-sub-source LR for the D12S391 locus then becomes 89.2053.

**Table 2.**
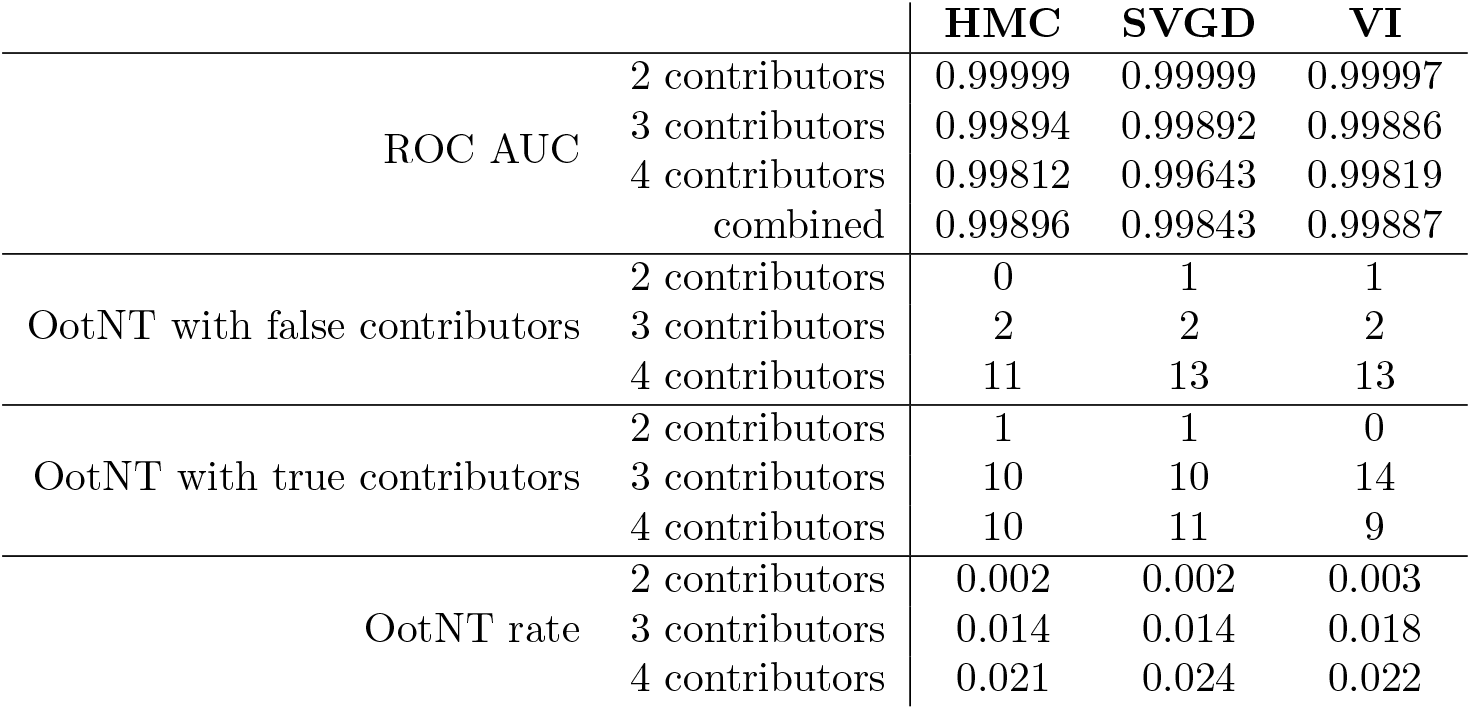
Performance metrics for the three tested algorithms (HMC: Hamiltonian Monte Carlo [15]; SVGD: Stein variational gradient descent [this work]; VI: variational inference with evidence lower-bound objective [this work]) on the ProvedIT benchmark [1]; OotNT = Opposite of the Neutral Threshold [14].

**Figure 5.**
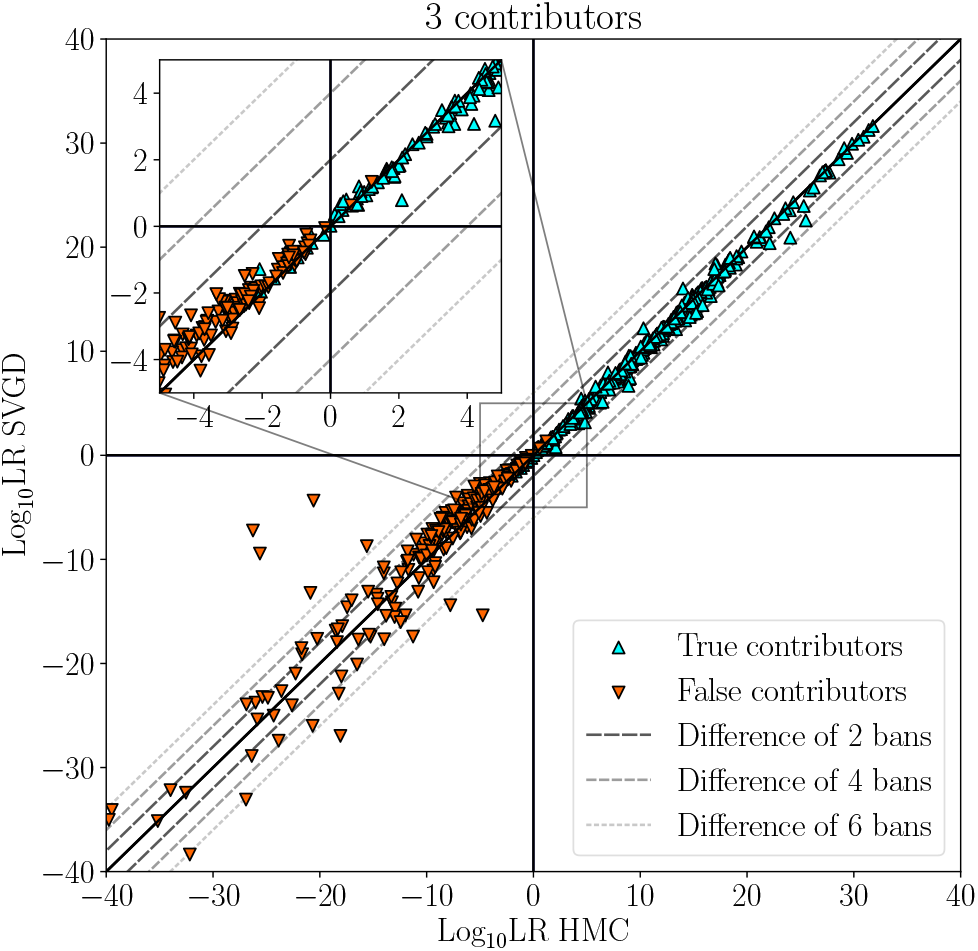
Comparison of results obtained by SVGD and HMC on 3-contributor mixtures.

**Figure 6.**
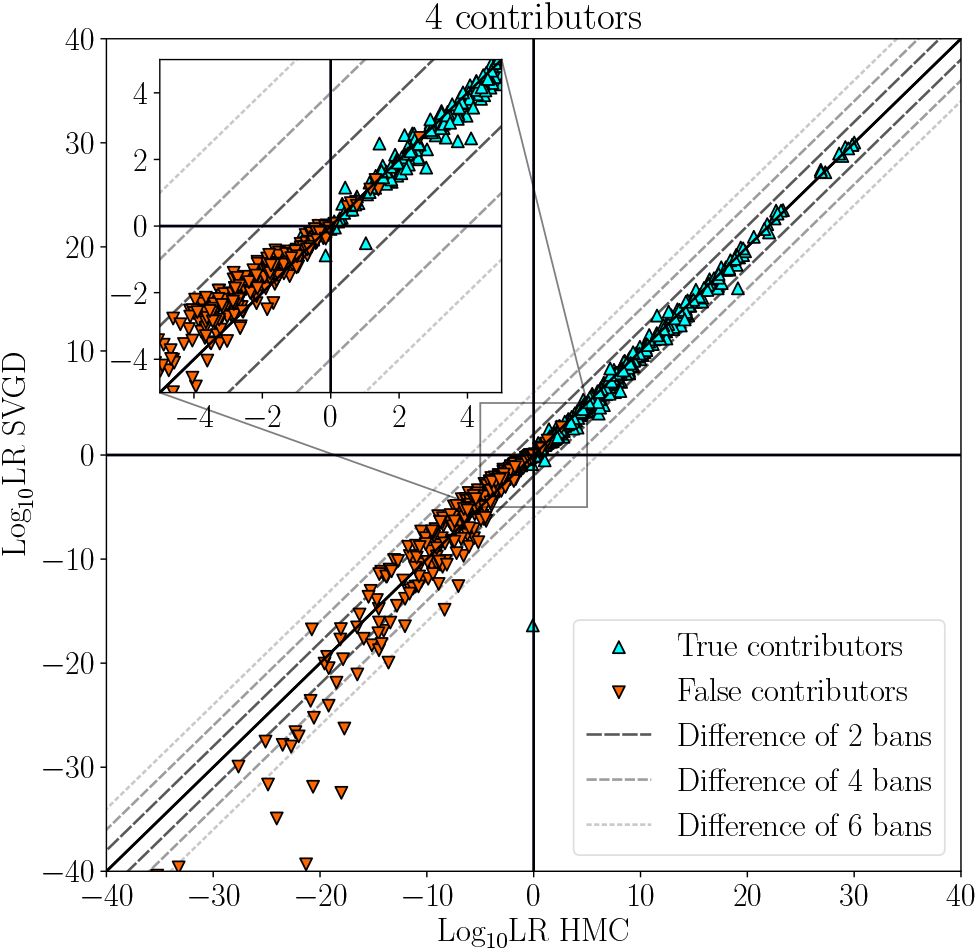
Comparison of results obtained by SVGD and HMC on 4-contributor mixtures. The scenario with log_10_ LR of -0.0852 in case of HMC and -16.3753 in case of SVGD is mixture E05 when Contributor 33 is assumed in the hypothesis of the prosecutor.

An interesting question is why SVGD is more sensitive to such missed peaks than both HMC and VI (log_10_ LR of 2.4072). It is known that SVGD is particularly prone to the curse of dimensionality and therefore tends to underestimate the variance of the posterior [2]. This is indeed seen in Figure 3, where SVGD underestimates (compared to the other estimators) the allele peak-height standard deviation. This increases the confidence of the model and, therefore, this estimator is less robust against extreme observations, such as an uncalled peak with a RFU larger than 1000.

Next, we consider the numbers of OotNT (Opposite of the Neutral Threshold [14]) scenarios with true and false contributors, as well as the OotNT rates [14]. The results are again given in Table 2. In these metrics, both SVGD and VI perform slightly worse than HMC. For example, VI provided 4 OotNT scenarios with true contribuors more than HMC for the 3-contributor mixtures, and both SVGD and VI provided 2 OotNT scenarios with false contribuors more than HMC for the 4-contributor mixtures. All of these scenarios are characterised by low certainty. For example, the additional 3-contributor OotNT scenarios with true contributors had log_10_ LRs of -0.7756, -0.2065, -0.2970, and -0.3269 in case of VI, and 0.1248, 0.3516, 0.2888, and 0.462 in case of HMC. The detailed results are available in Supplementary Material 1.

Visualisations of the full results are given in Figures 4 to 9. All comparisons confirm the strong agreement between the compared methods, with the difference between the log_10_ LRs from the different methods rarely exceeding 2 bans. In the few scenarios where differences are larger, both methods provide strong evidence. The results for HMC are taken from the supplementary materials of Ref. [14], where the priors of Riman et al. [12] were used, whereas here we use the modified priors described in Section 2.1. This indicates that the exact choice of priors is not crucial to the reported results.

### 3.2 Precision

A desirable property of a PG software is low run-to-run variability. It has been shown that HMC greatly reduces this variability compared to random-walk MCMC [15]. Low run-to-run variability implies high reproducibility of the results, which is known as *precision*. We compare the precision of HMC as previously benchmarked [14] with the precision of SVGD and VI. For this, we perform 10 independent repetitions of the analyses for each scenario and compare two statistics: the standard deviation of the resulting log_10_ LR and the difference between the largest and smallest log_10_ LR. The measurements are presented in Figure 10. SVGD is overall less precise than the other two algorithms, with 8 scenarios having a run-to-run standard deviation of the log_10_ LR larger than 0.2. The other two algorithms result in 3 scenarios each where the standard deviations are larger than 0.1. In 14 (HMC) and 13 (VI) scenarios, the standard deviations across runs are larger than 0.05. The precision of SVGD can be improved by either increasing the number of particles or using more iterations with learning-rate annealing. As we will see below, however, this would not compare well against VI in terms of the computational cost and is therefore not explored.

**Figure 7.**
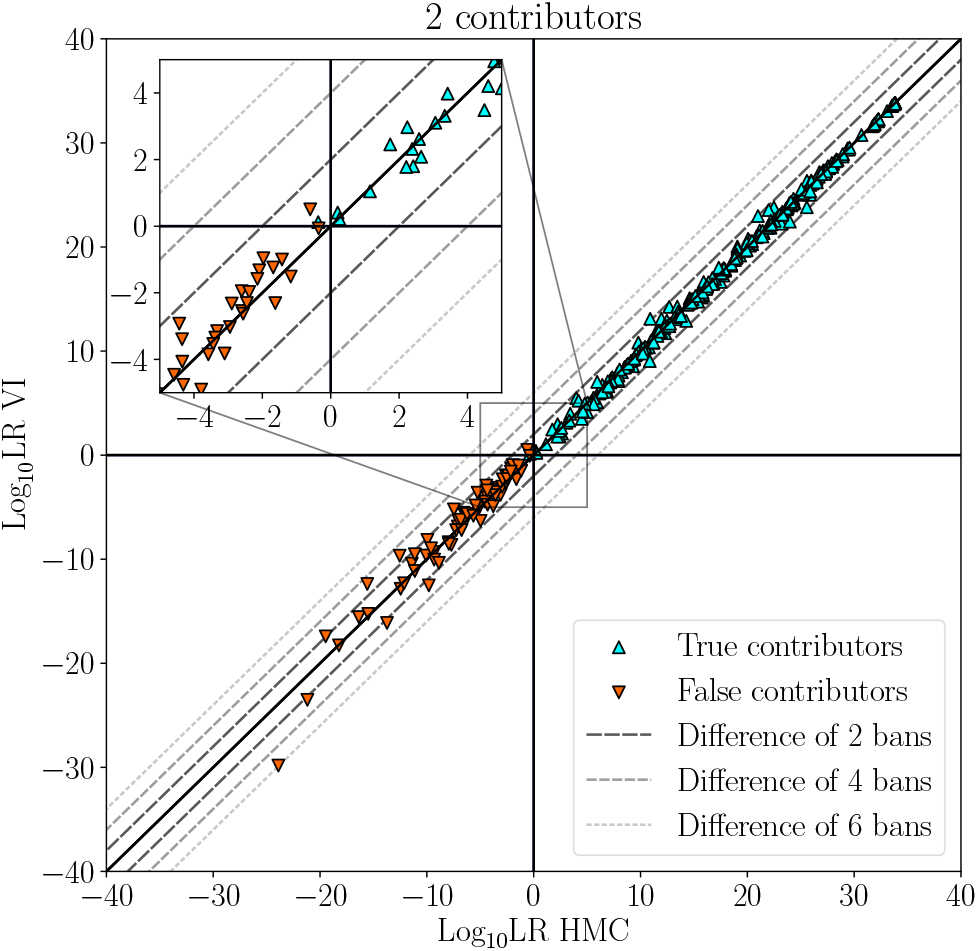
Comparison of results obtained by VI and HMC on 2-contributor mixtures.

**Figure 8.**
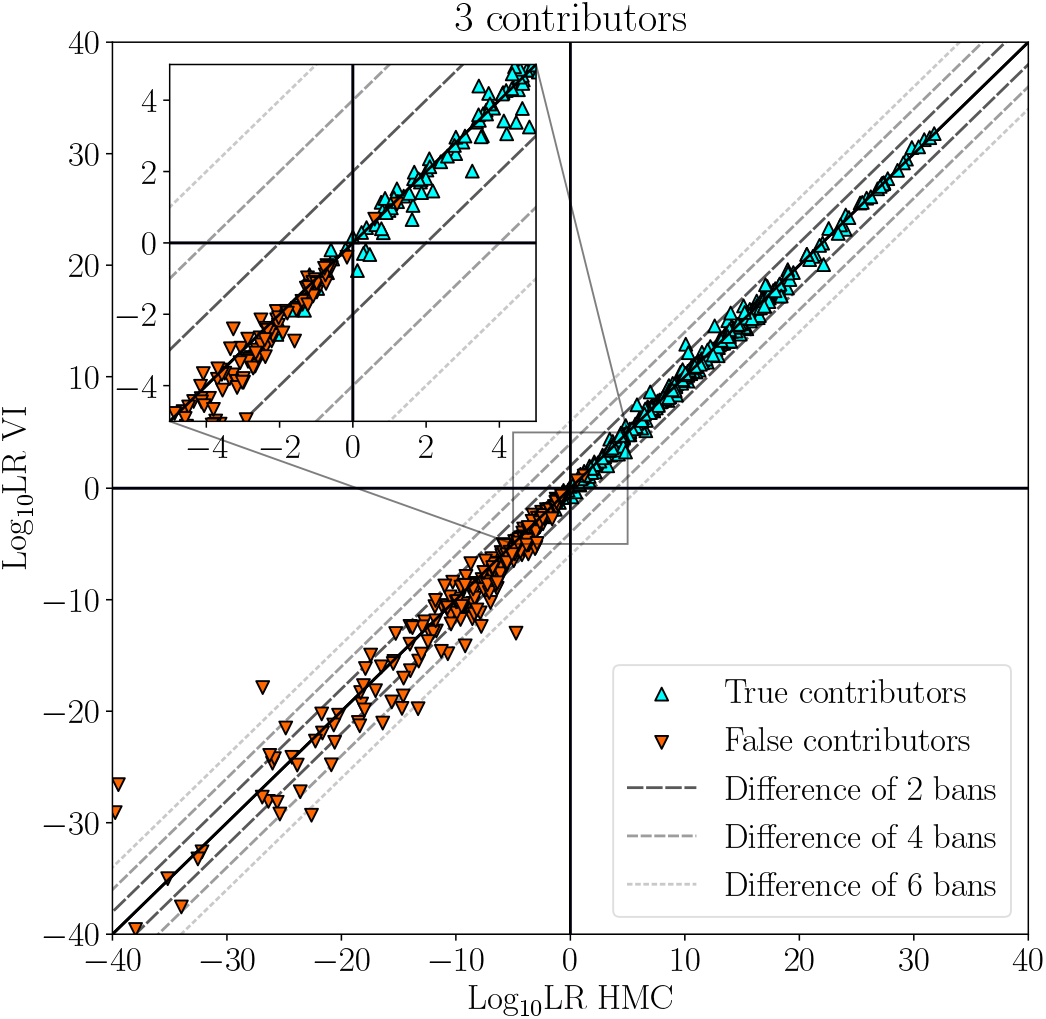
Comparison of results obtained by VI and HMC on 3-contributor mixtures.

**Figure 9.**
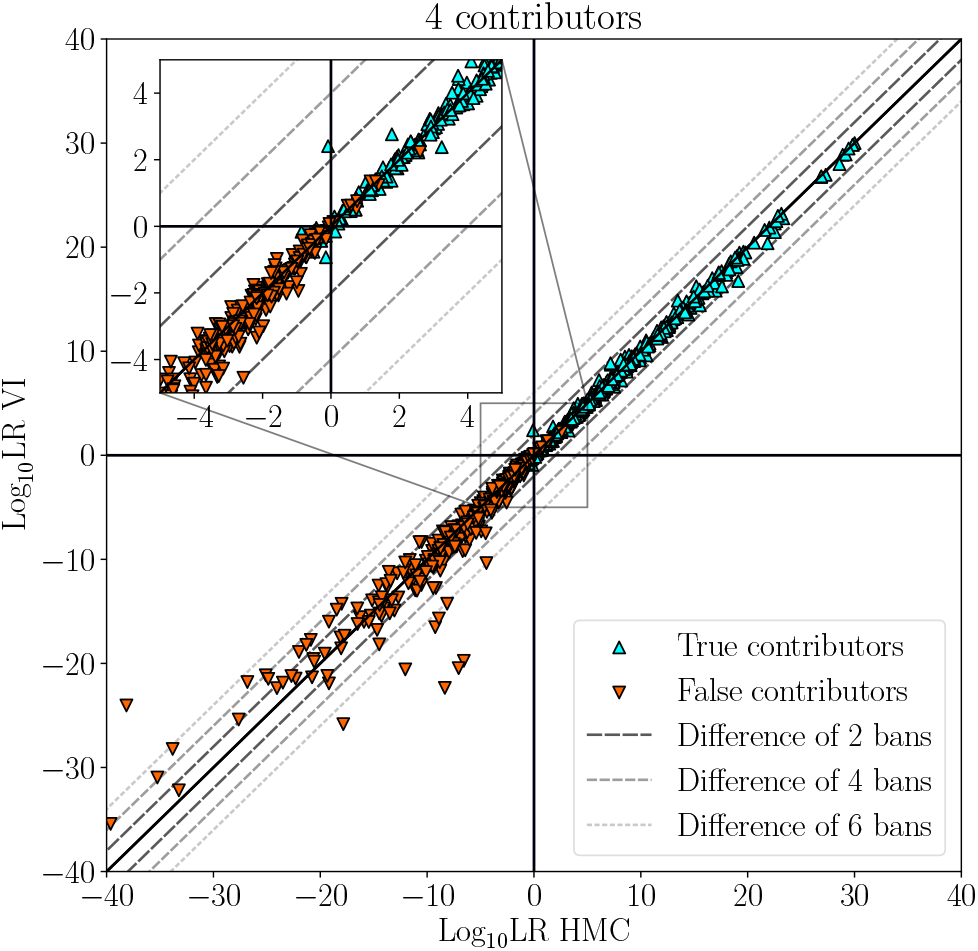
Comparison of results obtained by VI and HMC on 4-contributor mixtures.

**Figure 10.**
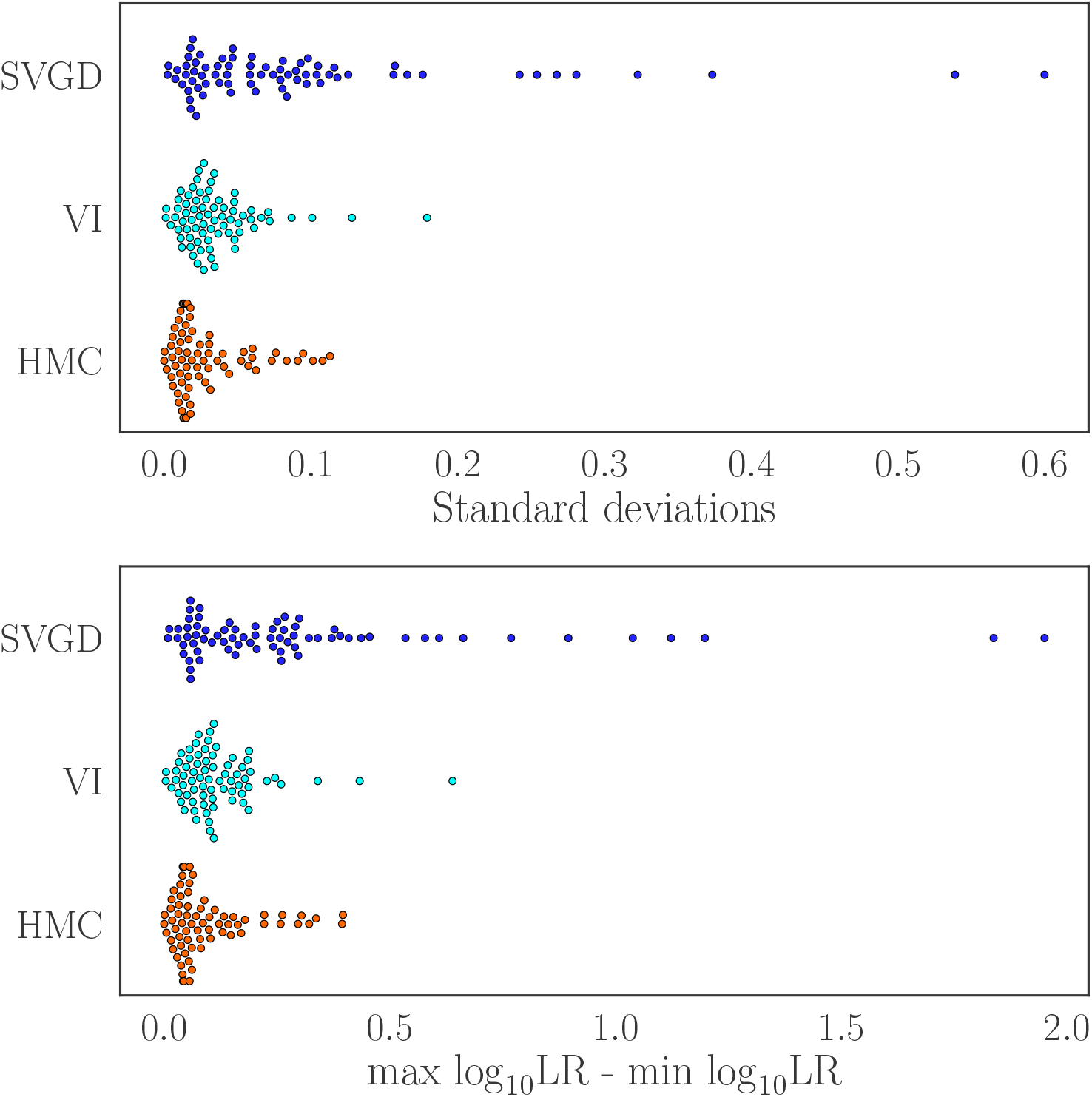
Precision comparison of the tested algorithms both in terms of the log_10_ LR standard deviation (top) and the difference between the largest and smallest log_10_ LR (bottom) across 10 independent repetitions for each scenario (1 scenario = 1 dot).

Visualisations of the full results are available in Supplementary Material 2, with the raw data provided in Supplementary Material 1.

### 3.3 Runtime

The third performance metric of interest is the computational runtime. Variational techniques are usually faster, which is the main reason why they became popular with the deep-learning community. We confirm this here for PG, observing significant speedups as shown in Figure 11. We compare the runtimes on identical computer hardware for all 4-contributor scenarios of the benchmark [12]. The computationally most demanding mixture takes 1 hour and 54 minutes to be solved with HMC. For SVGD, the longest observed inference time is 36 minutes 28 seconds. VI provides the largest speedups, with the slowest inference completing in 18 minutes and 27 seconds. On average, VI is 4.33 times faster than HMC and 1.72 times faster than SVGD on 4- contributor scenarios. In 81.1% of the 4-contributor scenarios (103/127) VI completes the inference in under 10 minutes and in 96.1% (122/127) in under 12 minutes. SVGD completes 91.4% (116/127) of these scenarios in under 20 minutes.

**Figure 11.**
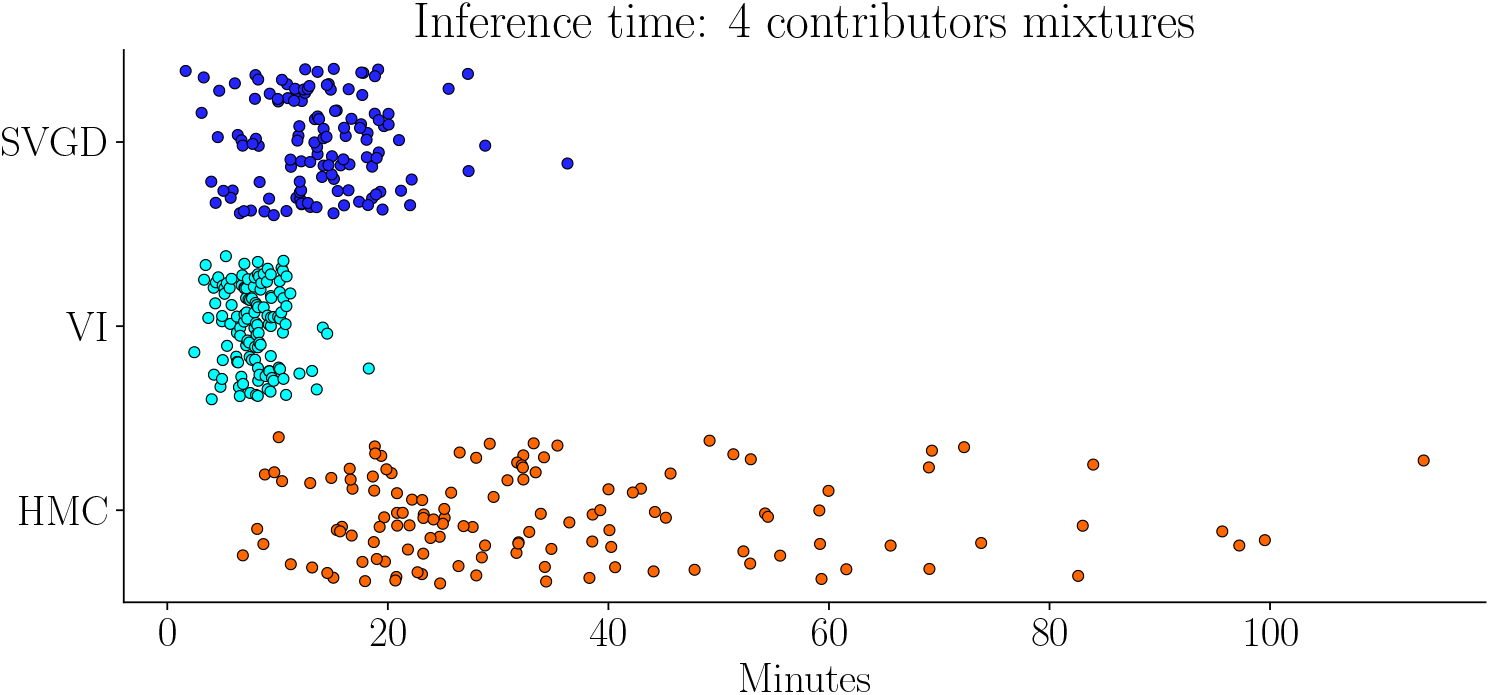
Comparison of the computational runtime of the different algorithms on the same 4-contributor mixtures. Each dot represents a different scenario.

These runtime benchmarks used GPU hardware. Most forensic laboratories, however, still use CPUs to compute. We therefore confirm the runtimes of the best-performing algorithm, VI, on four vCPU cores (AMD EPYC 7V12 Rome) for the F03 RD14−0003−48 49 50 29−1;4;4;4−M2U15−0.403GF−Q1.3 06.15sec mixture. For this scenario, VI runs for 10 minutes and 13 seconds on the GPU, whereas the same inference on the CPU takes 67 minutes and 14 seconds. We note, however, that the code contains no CPU-specific optimisations so that this figure could probably be improved if needed.

## 4 Conclusions

We presented two variational Bayesian inference algorithms for probabilistic genotyping (PG): Stein variational gradient descent (SVGD) and variational inference with an evidence lower-bound objective (VI). These are applicable to PG models that are free of singularities. We have therefore also shown how to adapt the PG models of STRmix™ [16] to allow for variational inference. We then described the algorithms and explained their working principles and underlying assumptions.

We then validated the algorithms and checked the validity of the assumptions on the ProvedIT mixtures from the NIST comparative study [12] and compared with the Hamiltonian Monte Carlo (HMC) method, which was recently benchmarked on the same data set [14]. All three methods, HMC, SVGD, and VI were comparable in terms of accuracy with HMC slightly more accurate than the other two. This could, however, also be due to the different priors used in the HMC model. Importantly, VI achieved significantly lower computational runtimes than both SVGD and HMC while maintaining the high precision of HMC. It therefore seems to offer the best trade-off between accuracy, precision, and runtime, suggesting that VI could replace MCMC-based algorithms in practice and provide better user experience due to faster runtimes. Faster runtimes would also enable laboratories to run independent repetitions of the inference in order to quantify reproducibility.

While not outperforming in the benchmarks, SVGD is an algorithmically interesting method that offers ample opportunity for optimization and links to established mathematical frameworks such as particle filters [4] for which efficient parallel software exists [5] While our implementation of SVGD was significantly faster than HMC, it was overly sensitive to missing peaks as seen in case of the E05 mixture.

An important limitation of the present work is that our implementation of VI was limited to a multivariate Gaussian approximation of the posterior. Recently, it has been shown that it is possible to construct more general approximations using VI, theoretically even a universal one [7]. This greater approximation power is achieved by learning invertible distribution transformations, called *normalising flows*. We tested possible flow architectures and obtained promising results with inverse autoregressive flows [10]. Then, we often observed improved quality of the posterior estimation, but the method had two significant drawbacks: First, the computation time increased to become comparable with that of HMC. Second, we could not find a robust way to prevent gradient divergence, which happened occasionally when normalising flows were used.

Taken together, our results not only provide a way of accelerating inference in Bayesian PG without sacrificing much accuracy or precision, but they also open the field of research toward a more diverse range of inference algorithms beyond sampling-based MCMC methods. We hope this might trigger a discussion in the field and reinvigorate the search for better, more scalable, and mathematically founded algorithms for DNA mixture deconvolution in forensic genetics.

## Acknowledgments

We thank Dr. Anne-Marie Pflugbeil and Holger Schönborn (both Qualitype GmbH, Dresden) for the support during the development of the project.

