## Supplementary Material 2 for "Variational inference accelerates accurate DNA mixture deconvolution"

October 2022

### 1 Visualisations of precision of the methods

In this document, we visualise the full precision benchmark data for the three compared methods for all scenarios in which all runs resulted in LR>0 for all methods. Each panel represents one scenario with 10 independent repetitions of the inference using the methods **SVGD**, **VI**, and **HMC** distinguished by colour. The scenarios are sorted lexicographically, and the y-axes show the same $\log_{10}$ LR ranges (i.e., same difference between upper and lower range bound) for visual comparability of the spread of the results.

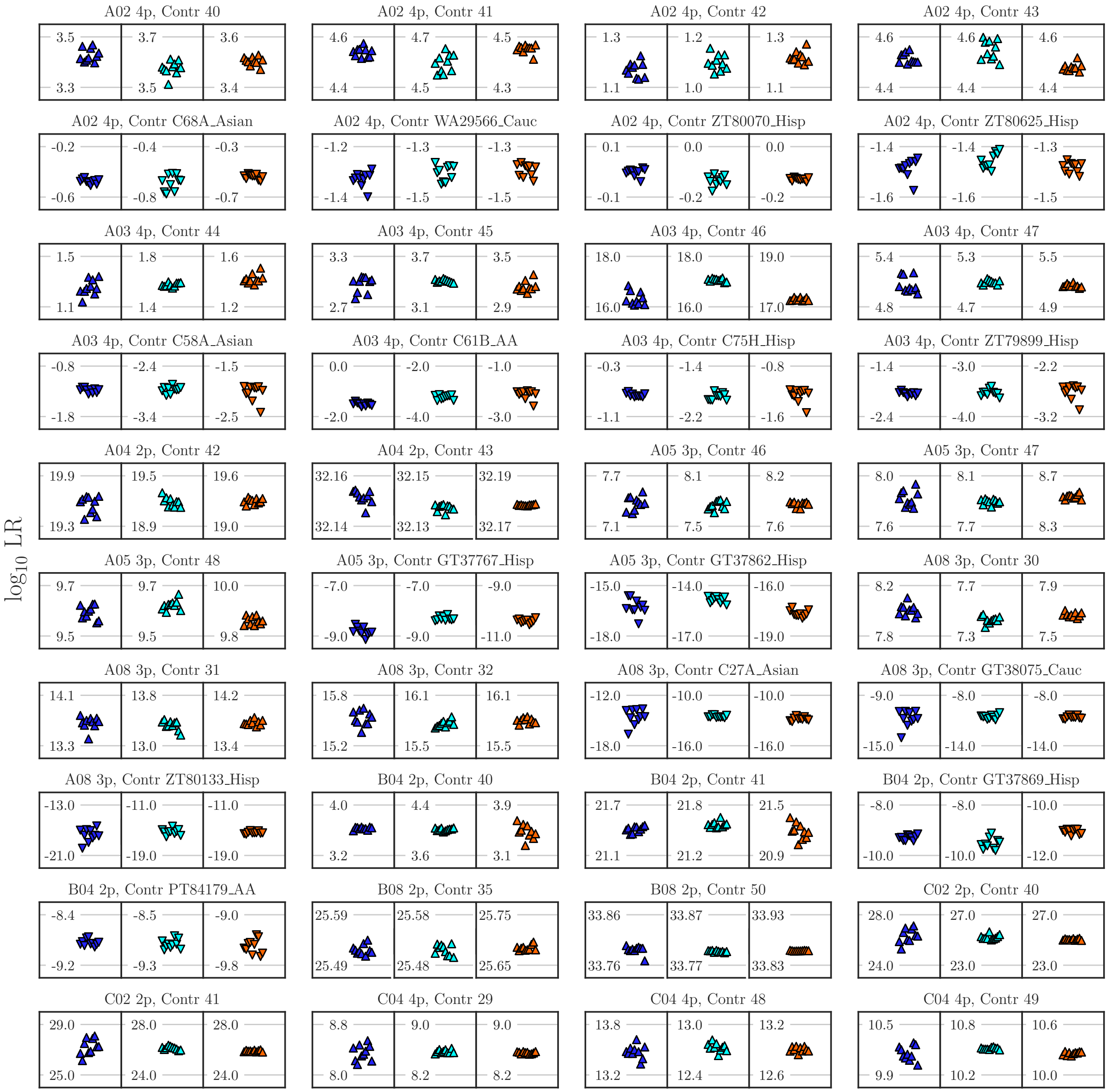

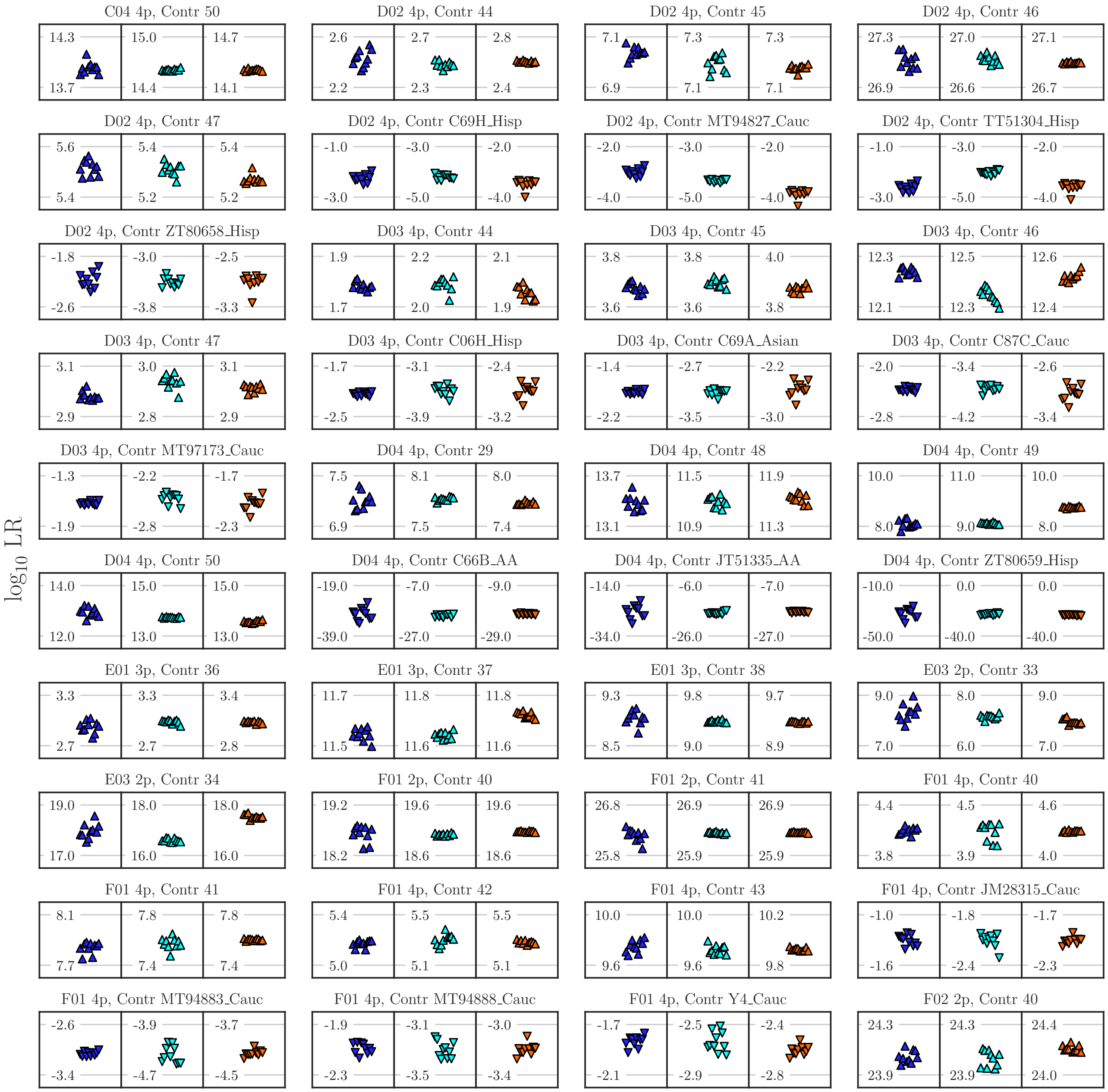
log10 LR
C04 4p, Contr 50
D02 4p, Contr 44
D02 4p, Contr 45
D02 4p, Contr 46
D02 4p, Contr 47
D02 4p, Contr C69H_Hisp
D02 4p, Contr MT94827_Cauc
D02 4p, Contr TT51304_Hisp
D02 4p, Contr ZT80658_Hisp
D03 4p, Contr 44
D03 4p, Contr 45
D03 4p, Contr 46
D03 4p, Contr 47
D03 4p, Contr C06H_Hisp
D03 4p, Contr C69A_Asian
D03 4p, Contr C87C_Cauc
D03 4p, Contr MT97173_Cauc
D04 4p, Contr 29
D04 4p, Contr 48
D04 4p, Contr 49
D04 4p, Contr 50
D04 4p, Contr C66B_AA
D04 4p, Contr JT51335_AA
D04 4p, Contr ZT80659_Hisp
E01 3p, Contr 36
E01 3p, Contr 37
E01 3p, Contr 38
E03 2p, Contr 33
E03 2p, Contr 34
F01 2p, Contr 40
F01 2p, Contr 41
F01 4p, Contr 40
F01 4p, Contr 41
F01 4p, Contr 42
F01 4p, Contr 43
F01 4p, Contr JM28315_Cauc
F01 4p, Contr MT94883_Cauc
F01 4p, Contr MT94888_Cauc
F01 4p, Contr Y4_Cauc
F02 2p, Contr 40

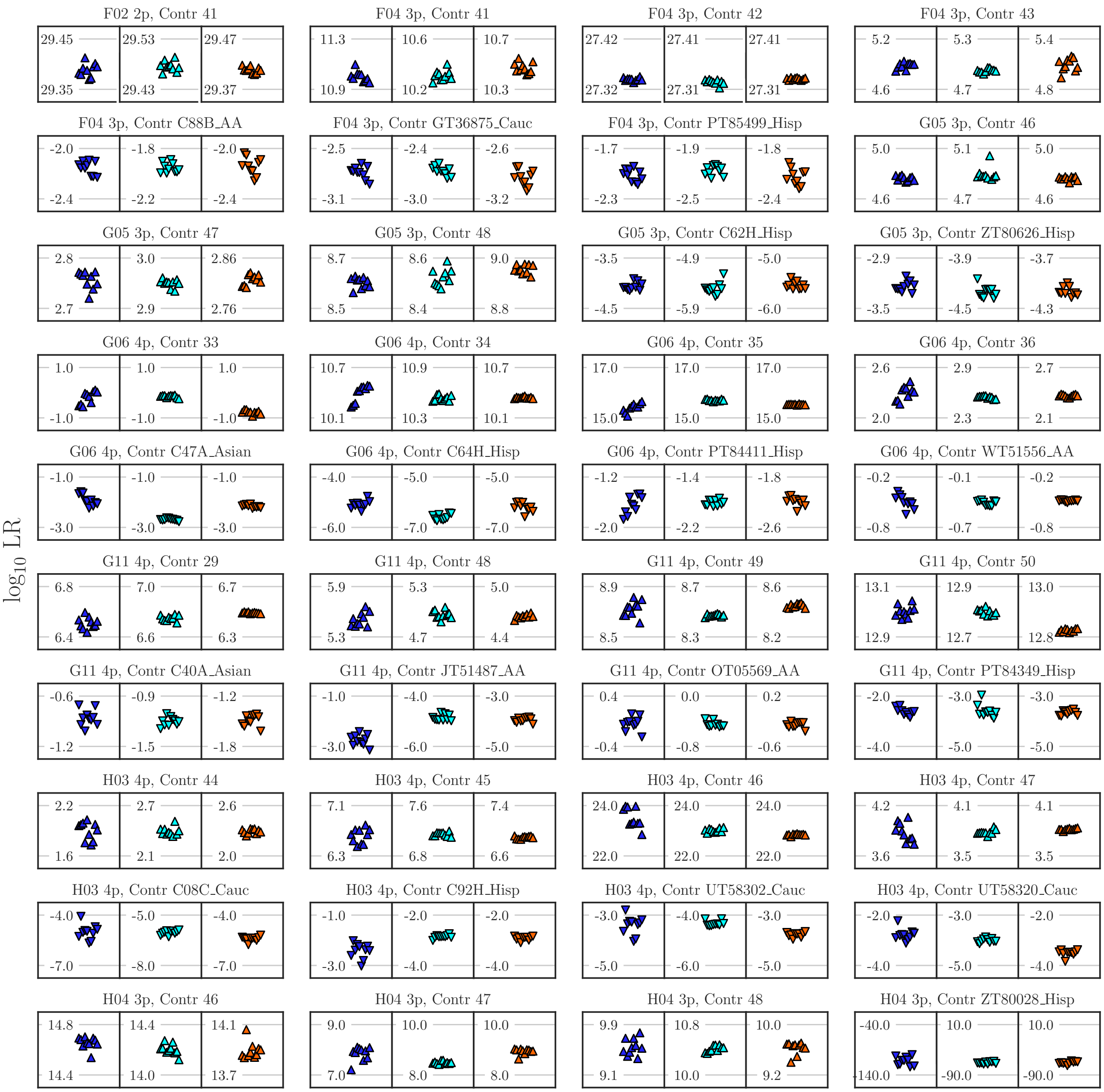
log10 LR
F02 2p, Contr 41
29.45 29.35 29.53 29.43 29.47 29.37
F04 3p, Contr 41
11.3 10.9 10.6 10.2 10.7 10.3
F04 3p, Contr 42
27.42 27.32 27.41 27.31 27.41 27.31
F04 3p, Contr 43
5.2 4.6 5.3 4.7 5.4 4.8
F04 3p, Contr C88B_AA
-2.0 -2.4 -1.8 -2.2 -2.0 -2.4
F04 3p, Contr GT36875_Cauc
-2.5 -3.1 -2.4 -3.0 -2.6 -3.2
F04 3p, Contr PT85499_Hisp
-1.7 -2.3 -1.9 -2.5 -1.8 -2.4
G05 3p, Contr 46
5.0 4.6 5.1 4.7 5.0 4.6
G05 3p, Contr 47
2.8 2.7 3.0 2.9 2.86 2.76
G05 3p, Contr 48
8.7 8.5 8.6 8.4 9.0 8.8
G05 3p, Contr C62H_Hisp
-3.5 -4.5 -4.9 -5.9 -5.0 -6.0
G05 3p, Contr ZT80626_Hisp
-2.9 -3.5 -3.9 -4.5 -3.7 -4.3
G06 4p, Contr 33
1.0 -1.0 1.0 -1.0 1.0 -1.0
G06 4p, Contr 34
10.7 10.1 10.9 10.3 10.7 10.1
G06 4p, Contr 35
17.0 15.0 17.0 15.0 17.0 15.0
G06 4p, Contr 36
2.6 2.0 2.9 2.3 2.7 2.1
G06 4p, Contr C47A_Asian
-1.0 -3.0 -1.0 -3.0 -1.0 -3.0
G06 4p, Contr C64H_Hisp
-4.0 -6.0 -5.0 -7.0 -5.0 -7.0
G06 4p, Contr PT84411_Hisp
-1.2 -2.0 -1.4 -2.2 -1.8 -2.6
G06 4p, Contr WT51556_AA
-0.2 -0.8 -0.1 -0.7 -0.2 -0.8
G11 4p, Contr 29
6.8 6.4 7.0 6.6 6.7 6.3
G11 4p, Contr 48
5.9 5.3 5.3 4.7 5.0 4.4
G11 4p, Contr 49
8.9 8.5 8.7 8.3 8.6 8.2
G11 4p, Contr 50
13.1 12.9 12.9 12.7 13.0 12.8
G11 4p, Contr C40A_Asian
-0.6 -1.2 -0.9 -1.5 -1.2 -1.8
G11 4p, Contr JT51487_AA
-1.0 -3.0 -4.0 -6.0 -3.0 -5.0
G11 4p, Contr OT05569_AA
0.4 -0.4 0.0 -0.8 0.2 -0.6
G11 4p, Contr PT84349_Hisp
-2.0 -4.0 -3.0 -5.0 -3.0 -5.0
H03 4p, Contr 44
2.2 1.6 2.7 2.1 2.6 2.0
H03 4p, Contr 45
7.1 6.3 7.6 6.8 7.4 6.6
H03 4p, Contr 46
24.0 22.0 24.0 22.0 24.0 22.0
H03 4p, Contr 47
4.2 3.6 4.1 3.5 4.1 3.5
H03 4p, Contr C08C_Cauc
-4.0 -7.0 -5.0 -8.0 -4.0 -7.0
H03 4p, Contr C92H_Hisp
-1.0 -3.0 -2.0 -4.0 -2.0 -4.0
H03 4p, Contr UT58302_Cauc
-3.0 -5.0 -4.0 -6.0 -3.0 -5.0
H03 4p, Contr UT58320_Cauc
-2.0 -4.0 -3.0 -5.0 -2.0 -4.0
H04 3p, Contr 46
14.8 14.4 14.4 14.0 14.1 13.7
H04 3p, Contr 47
9.0 7.0 10.0 8.0 10.0 8.0
H04 3p, Contr 48
9.9 9.1 10.8 10.0 10.0 9.2
H04 3p, Contr ZT80028_Hisp
-40.0 -140.0 10.0 -90.0 10.0 -90.0
